# GABA and Glutamate response to social processing; a functional MRS study

**DOI:** 10.1101/2023.05.17.541109

**Authors:** Duanghathai Pasanta, David J. White, Jason L. He, Talitha C. Ford, Nicolaas A. Puts

**Affiliations:** Department of Forensic and Neurodevelopmental Sciences, Sackler Institute for Translational Neurodevelopment, Institute of Psychiatry, Psychology, and Neuroscience, King’s College London, London, United Kingdom; Department of Radiologic Technology, Faculty of Associated Medical Sciences, Chiang Mai University, Chiang Mai, Thailand; Centre for Human Psychopharmacology & Swinburne Neuroimaging, School of Health Sciences, Swinburne University of Technology, Melbourne, Australia; MRC Centre for Neurodevelopmental Disorders, King’s College London, London, United Kingdom; Cognitive Neuroscience Unit, Faculty of Health, Deakin University, Geelong, Australia

## Abstract

Several studies have suggested that atypical social processing in multiple psychiatric conditions (e.g., autism) is associated with differences in excitation and inhibition, through changes in the levels of glutamate and GABA levels. While associations between baseline metabolite levels and behaviours can be insightful, assessing the neurometabolic response of GABA and Glutamate during social processing may inform altered neurochemical function in more depth. Thus far, there have been no attempts to determine whether changes in metabolite levels are detectable using functional magnetic resonance spectroscopy (fMRS) during social processing in a control population. We performed MEGA-PRESS edited fMRS to measure the dynamic response of GABA and glutamate in the superior temporal sulcus (STS) and visual cortex (V1) while viewing social stimuli, using a design that allows for analysis in both block and event-related approaches. Sliding window analyses were used to investigate GABA and glutamate dynamics at higher temporal resolution. A small decrease in GABA levels was observed during social stimulus presentation in V1, but no change was observed in STS. Conversely, non-social stimulus elicited changes in both GABA and glutamate levels in both regions. We discuss the feasibility of using fMRS analysis approaches to assess changes in metabolite response during social processing.

## Introduction

Social processing refers to the perception and interpretation of self and others and their behavioral responses, which result from interpersonal (social) interactions^1^. Social processing also covers facial emotion recognition, body language comprehension, and the ability to predict and infer the mental state of others from these non-verbal social clues^2^. Several studies have suggested that social processing differences observed in multiple psychiatric conditions (e.g., autism spectrum disorder (ASD)^3, 4^, schizophrenia^5, 6^, and anxiety disorder^7^) are associated with an aberrant excitatory-inhibitory (E/I) neurotransmission ^8–10^.

It is generally acknowledged that proton magnetic resonance spectroscopy (^1^H-MRS) is the only neuroimaging technique that allows for non-invasive in vivo quantification of neurometabolites within pre-defined regions of the brain^11^. ^1^H-MRS is a well-established technique for robust measurement of multiple neurometabolites in a single acquisition and has been extensively applied in both non-clinical and clinical groups^12, 13^. These MRS-measured metabolites can be directly linked to brain function and responses including excitatory and inhibitory tone^14, 15^. Two metabolites that are of particular interest are the primary excitatory neurotransmitter glutamate and the primary inhibitory neurotransmitter gamma-aminobutyric acid (GABA).

While the vast majority of MRS studies of glutamate and GABA have been performed at rest, being able to determine dynamic changes in glutamate/GABA balance is of increased interest to research in health and disease. MRS of glutamate and GABA (and particularly that of the latter) requires tailored techniques due to its low concentrations in the brain (and thus, low Signal to Noise; SNR) and overlap of more highly concentrated metabolites. J-difference editing is mostly typically used to measure GABA, and provides simultaneous measures of glutamate and glutamine (Glx). Despite their low concentration and overlap with more highly concentrated metabolites, recent developments in ^1^H-MRS techniques and MR instruments have led to increased temporal resolution and robust quantification of glutamate and GABA^16^.

In an attempt to capture dynamic metabolic changes due to stimulation^2^, MRS data acquired across completion of a task paradigm can be probed for metabolite fluctuations within windows of varying task demands. This emerging technique is called functional MRS (fMRS)^3, 24^. Recent J-difference editing fMRS works have demonstrated the feasibility of measuring activation-induced neurometabolite responses during various types of physiological stimulation, including in visual, motor, cognitive and pain domains^18, 19^, with an effective temporal resolution as low as 12 seconds^19, 20^. fMRS is of particularly interest for neurodevelopmental conditions where E/I balance is thought to be shifted, such as in autism or schizophrenia, since it enables the study of GABA and glutamate change directly in response to a stimulus. However, higher temporal resolution comes at a cost of lower SNR from low number of transients used compared to typical MRS measurement. Two main types of fMRS paradigms are used to probe metabolite fluctuations during stimulation: block and event-related designs. Block design paradigms hold the advantage of robust metabolite quantification by averaging over a large number of stimuli/trials and thus high SNR but require long fMRS measurements in a block. By contrast, in event-related designs, stimuli are time-locked with the fMRS acquisition and therefore more directly reflect the brain response to the stimulus, however this comes at the cost of low SNR. Thus, both approaches have their pro’s and con’s (described in more detail here^19, 21^).

With the ultimate goal of investigating altered E/I balance and its relation to social functioning in autism, we are first interested in investigating whether fMRS can be used to assess metabolic changes during social processing in a non-clinical adult population. Thus far, there have been no attempts to determine whether changes in metabolite levels are detectable using fMRS during social processing. Here, we performed fMRS in non-clinical young adults to assess the temporal resolution of GABA+ (GABA + macromolecules) and Glx (glutamate + glutamine) response during social processing *in-vivo*, with a paradigm allowing for assessment in both block and event-related responses. We hypothesized that that each stimulus presentation within each block affects metabolite levels differently. We measure metabolite dynamics in two brain regions: right superior temporal sulcus (STS), a region considered to play a major role in social perception and cognition^22^; and the occipital cortex containing V1, an early visual processing brain region (V1) that is not associated with social processing. We subsequently explore both block/event-related analyses as well as dynamic sliding window and change-point analyses to determine best analytical approaches to observe relevant changes, accounting for potential metabolite lag after stimulus presentation.

## Methods

### Participants

All studies were performed in accordance with the procedures approved by the Swinburne’s Human Research Ethics Committee (SUHREC) in line with the National Statement on Ethical Conduct in Human Research. Data were acquired from non-clinical adults aged 18-40 years. Participants were free from psychiatric, genetic, and neurological conditions, medications, nicotine and caffeine (abstained 24 hours), and recreational drugs (abstained 1 week). All data were acquired after written informed consent was obtained.

### MR data acquisition

Magnetic resonance imaging (MRI) and MRS data were acquired using a Siemens Trio 3T MRI scanner (Siemens, Erlangen, Germany) with 32-channel head coil at Swinburne University of Technology, Melbourne, Australia. T1-weighted structural images for MRS voxel localization were acquired using a magnetisation pre-prepared rapid gradient echo (MPRage) pulse sequence with an inversion recovery (176 slices, slice thickness =1.0mm, voxel resolution = 1 mm^3^, TR = 1900 ms, TE = 2.52 ms, TI = 900ms, flip angle = 9°, field of view 350mm × 263mm × 350 mm, acquisition time = 5min).

After structural image acquisition, fMRS was performed using the Mescher-Garwood Point Resolved Spectroscopy (MEGA-PRESS) sequence^23^ (TE/TR 68/2000 ms, 2048 data points) with editing pulses placed at 1.9 (edit-ON) and 4.8 (edit-OFF) ppm. Voxels were placed in the right STS (4 × 3 × 2 cm^3^), and bilateral V1 (3 × 3 × 3 cm^3^). During MRS data acquisition, participants were presented with four stimulus blocks: 1) resting MRS (fixation cross, 256 transients), 2) fMRS (dynamic faces or objects, 200 transients), 3) fMRS resting MRS (fixation cross, 200 transients), and 4) fMRS (dynamic faces or objects, 200 transients, order counterbalanced). The four stimulus blocks were repeated for both STS and V1 voxels (Figure 1).

**Figure 1.**
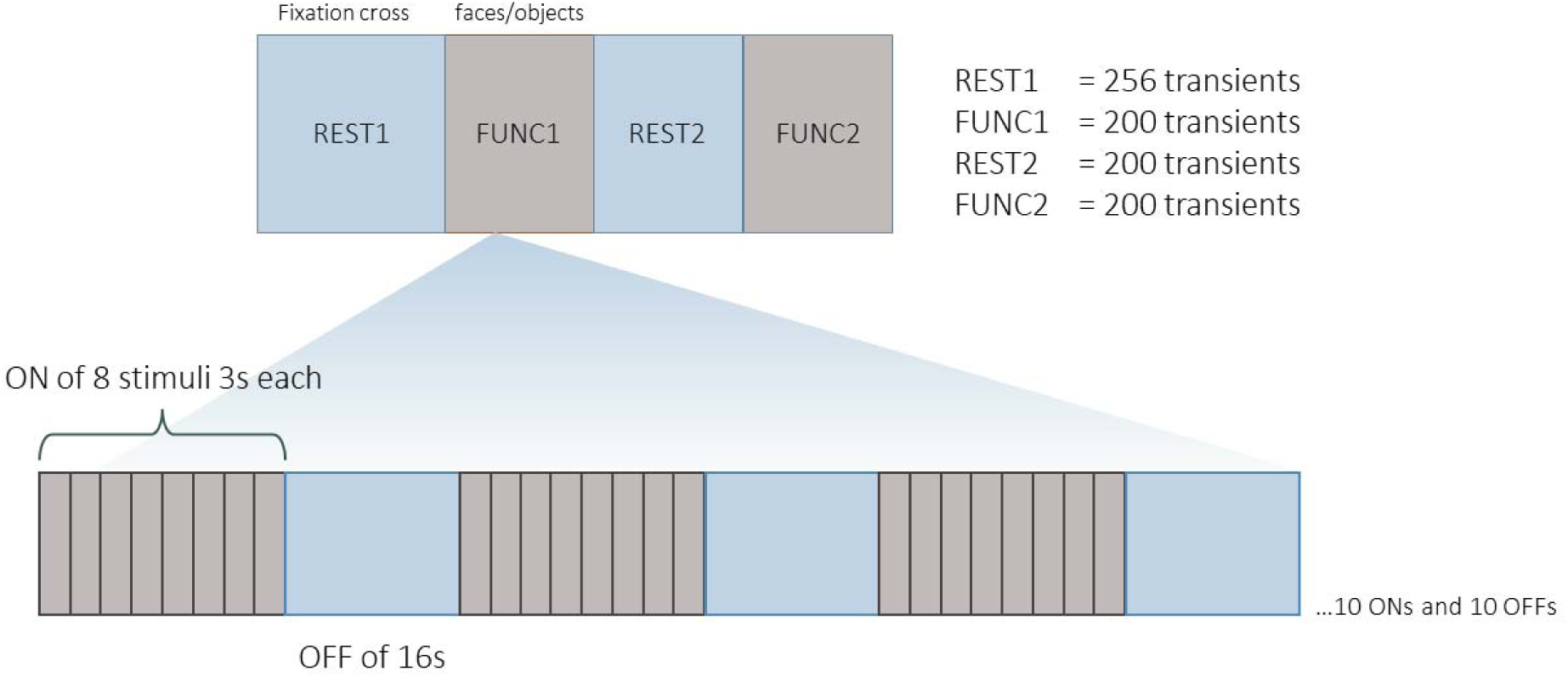
fMRS experimental design. A total of four MRS acquisition blocks were performed. During functional blocks (FUNC1 and FUNC2), participants were asked to passively watch 3s movie clips of dynamic faces or dynamic objects (24 s stimulus-ON, 16 s stimulus OFF) (order counterbalanced).

Within each functional block (second and fourth blocks), social (faces) and non-social (objects) stimuli were presented as sub-blocks (24 sec stimulus-ON, 16 sec stimulus-OFF), making these data a combined block and event-related design. A recent fMRI study has shown that the dynamic face stimuli used in this study reliably activates the STS, a central social brain region^22^, while the objects do not. Stimuli were 3 sec videoclips of dynamic faces or dynamic objects. There were eight 3-second videos presented in each sub-block, and there were a total of 10 subblocks for each functional stimulus block (see Figure 1). Total time for each functional block was 6 min 40 sec.

### Data analysis

#### MRS data analysis

Using an in-house modified version of Gannet^24^, data were pre-processed^25^ and fit using standard frequency and phase correction and fitting approaches within-block and expressed relative to internal reference total creatine (creatine + phosphocreatine: tCr)^26^. Using total Cr (tCr) as internal reference allows for minimizing influence of changes during the data acquisition (e.g., scanner drift, change in linewidth, chemical shift displacement)^27^, which is beneficial to a within-subject design. Glutamine and glutamate at 2.2 – 2.4 ppm were quantified as Glx. We should note that with MEGA-PRESS, the GABA signal at 3ppm is co-edited with macromolecule signal, and hence will be referred to as GABA+. MRS quantification therefore yielded metabolite levels corresponding to each voxel location relative to tCr, for example GABA+/tCr and Glx/tCr. Schematic representations of analyses performed is in Figure 2. The analysis and quantification scripts can be found at the Open Science Framework at https://osf.io/y7jqp/.

**Figure 2.**
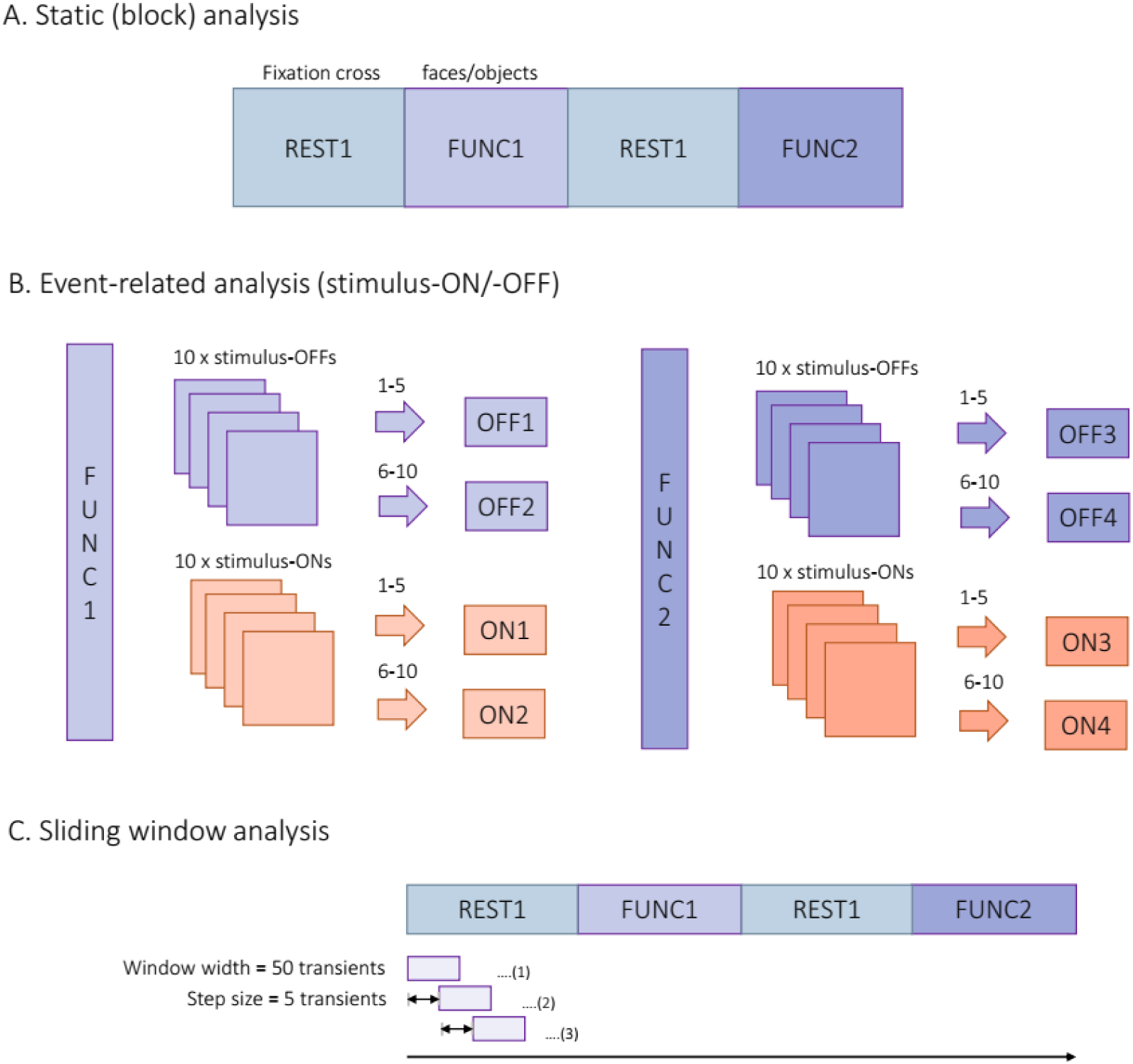
(A) Static (block) analysis based on stimulus paradigm of rest blocks and functional blocks. (B) Event-related analysis based on stimulus presentation TRs within each functional block. The stimulus ON and OFF periods were divided into ON1, ON2, OFF1, and OFF2 for FUNC1, and ON3, ON4, OFF3, and OFF4 for FUNC2. (C) Sliding window analysis with window width of 50 transients and step size of 5 transients.

##### Static window and event-related analysis

Data were analysed in blocks of rest (REST1 and REST2) and functional (FUNC1 and FUNC2). Due to the nature of block analysis, we did not separate for TRs corresponding to stimulus type. Because each functional block contains both stimulus-ON and stimulus-OFF blocks, we further analysed differences within-block by comparing TR periods corresponding to stimulus-ONs (24 sec stimulus-ONs of 3 sec movie clips of dynamic faces or dynamic objects) and stimulus-OFFs (16 sec rest), allowing for event-related analysis ^28^. We then halved the stimulus-ON and stimulus-OFF for each FUNC block and REST block to gain sufficient SNR for MRS spectral fitting (ON1, ON2, OFF1, OFF2 for FUNC1; ON3, ON4, OFF3, OFF4 for FUNC2).

##### Sliding window analysis

Metabolite levels were calculated by averaging the spectra over a window width of 50 transients, with a step size of 5 transients, regardless of the stimulus type (faces or objects). This allows for creating effective time course of metabolite levels as they change through time, with an effective temporal resolution of 1 minute and 40 seconds for 50 transients and effective time course of 10 seconds for 5 transients.

#### Statistics

Data were analysed in R (Version 4.1.1). Data were first screened for data skewness/kurtosis using Shapiro-Wilk tests. Outliers were removed based on the absolute deviation from the median of each metabolite with a threshold of 2.5 ^29, 30^.

##### Blocks (static) analysis

Data between averaged spectra blocks (static) were compared for changes in metabolite levels across stimulus blocks (RESTs or FUNCs) using one-way ANOVA test. Non-parametric statistic test of Kruskal-Wallis was used in subsequent analysis to compare metabolite levels across stimulus blocks with different stimulus types (faces or objects) due to data skewness.

##### Event-related analysis

Linear mixed-effects models was performed using the R package *lmerTest*^31^ to assess the relationship between metabolite levels and stimulus-ON/stimulus-OFF within functional blocks, and to investigate whether there was a possible lag in metabolite responses within the functional blocks (i.e., response delay and bleed into subsequence stimulus-OFFs). The sub-blocks of stimulus-ONs or stimulus-OFFs of each functional block were added as the fixed effect. The participants were included as intercept for random effect in the model. This analysis was done irrespective of stimulus types.

##### Sliding window analysis

For sliding window analysis, a Linear Mixed-Effects Model^31^ was used to investigate the effect of stimulus type and stimulus block on metabolite levels. The new variables to represent the combination of session and stimulus type were used as fixed effects, i.e., REST1, FUNC1+faces, FUNC1+objects, REST2, FUNC2+faces and FUNC2+objects. The time of measurement from the start of the experiment were included as fixed effects along with their interaction, and participants were included as a random effect.

A changepoint analysis (R *mcp*^32^ package) was performed on sliding window data to detect when metabolite levels first changed in pattern. The mean metabolite levels were calculated for each timepoint. A piecewise regression model was used with an autoregressive (AR) component to accounted for autocorrelation between datapoints. The model had two segments, each representing a different relationship between the metabolite and time, with a change point separating the two segments. In the first segment, the metabolite levels were modelled as a constant and an AR (1), to model the dependency of current value on its previous value. In the second segment, the metabolite levels were modelled as a function of time and an AR (1) process, but without an intercept. The resulting plots from *mcp* changepoint analysis showed the posterior distribution of the change point location based on a Bayesian inference approach^32–34^. The spread of the posterior distribution reflects uncertainly or the variability of the changepoint in the data, where the total probability is equal to 1 ^32^.

## Results

Data were acquired from 11 non-clinical adults aged between 18-40 years (mean age 26.5 years old; 6 females). Total of 8 spectra were acquired from each participant for either rest or functional block (V1 n=4, STS n=4). Table 1 shows quality metric for each analysis. Figure 3 shows example spectra from both static window analysis and sliding window analysis from both V1 and STS region during FUNC1.

**Figure 3.**
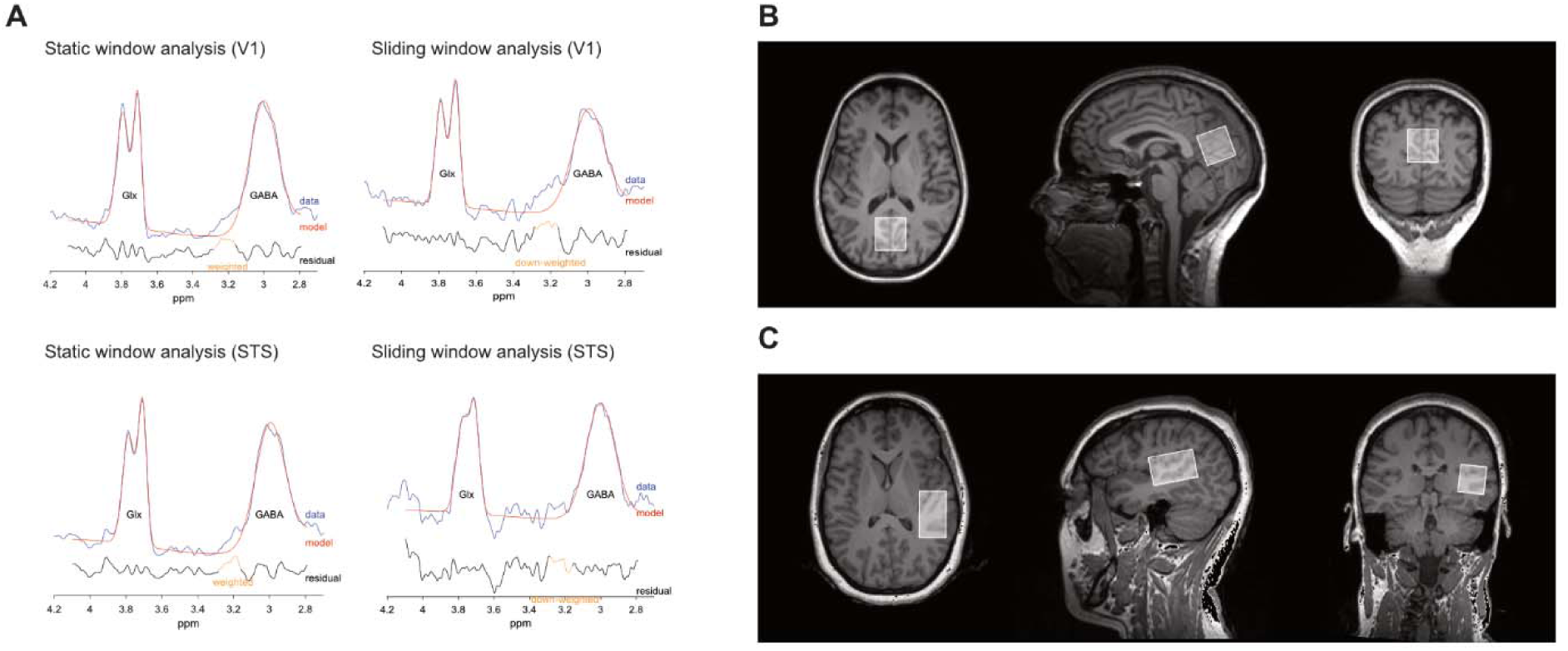
(A) Example spectra from static (block) window analysis (number of transients = 200) and sliding window analysis (number of transients = 50) during FUNC1. (B) Example voxel placement on V1 region and (C) STS region.

**Table 1.** quality metric for each analysis approach.

| Analysis approach | Brain region | NAA FWHM | NAA SNR | GABA+ fit error | Glx fit error |
| --- | --- | --- | --- | --- | --- |
| Static | V1 | 8.22 (0.77) | 428.19<br>(90.21) | 2.65 (0.44) | 2.57 (0.54) |
|  | STS | 15.00 (3.68) | 293.89<br>(59.01) | 2.93 (0.56) | 2.94 (0.64) |
| Event-related/ON | V1 | 8.19 (0.78) | 228.18<br>(40.83) | 4.16 (0.77) | 4.03 (0.83) |
|  | STS | 14.97 (3.72) | 161.58<br>(31.44) | 4.8 (0.99) | 4.67 (1.16) |
| Event-related/OFF | V1 | 8.21 (0.81) | 183.66<br>(30.83) | 4.55 (0.78) | 4.37 (0.93) |
|  | STS | 14.87 (3.51) | 124.95<br>(29.59) | 6.04 (1.38) | 5.57 (1.69) |
| Sliding window | V1 | 8.22 (0.77) | 214.26<br>(37.46) | 4.29 (0.84) | 4.17 (0.99) |
|  | STS | 15.02 (3.66) | 148.38<br>(31.01) | 5.23 (1.19) | 5.04 (1.32) |

### Static window analysis

Based on static (block) level analysis (see Figure 4), we estimated GABA+/tCr and Glx/tCr ratios across both stimulus types combined and found no significant changes between functional and rest blocks (p > 0.1). Subsequent analyses based on stimulus type also found no significant change in both GABA+/tCr and Glx/tCr for either faces or objects stimuli (p > 0.1) (Supplementary Figure 1 and Supplementary Figure 2).

**Figure 4.**
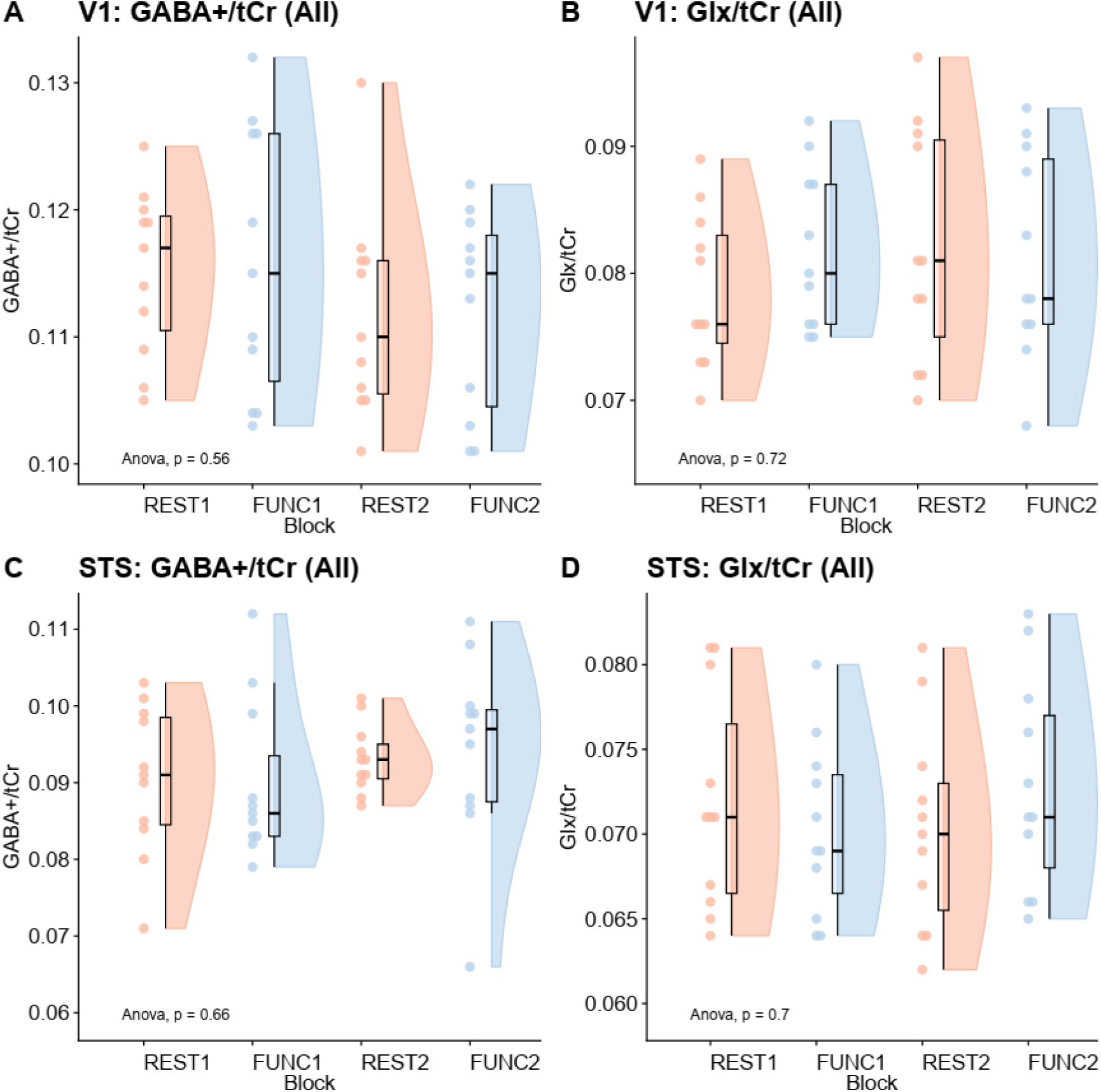
GABA+/tCr and Glx/tCr levels in (A, B) V1 and (C, D) STS region based on functional magnetic resonance spectroscopy (fMRS) blocks across both functional stimulus types (i.e., faces and objects). FUNC1, FUNC2: participants passively viewed social (faces) or non-social (objects) stimuli, counterbalanced; REST1, REST2: participants passively viewed fixation cross.

We then further investigated event-related changes by comparing metabolite levels from transients corresponding to stimulus-ON and stimulus-OFF TRs separately, regardless of stimulus type. Figure 5 illustrates GABA+/tCr and Glx/tCr levels across stimulus-ON and stimulus-OFF in STS. No statistical significances in metabolite measures were found in the STS region (p > 0.1). Figure 6 shows the GABA+/tCr and Glx/tCr levels across stimulus-ON and stimulus-OFF in V1. The fixed effects model of Glx/tCr in V1 shows that for stimulus-OFF, Glx/tCr increased compared to baseline (OFF1) during the second half of the first functional block (FUNC1) when the stimulus was OFF (OFF2) and during the first half of FUNC2 when stimulus was off (OFF3) (OFF2: β = 0.005, *SE* = 0.002, t = 2.36, p = 0.02, OFF3: β = 0.005, *SE* = 0.002, t = 2.36, p = 0.0016). For the fixed effects model for GABA+/tCr in V1, the result showed a decrease during the second half of stimulus-ON in the second functional block (FUNC2, ON4) (β = −0.009, *SE* = 0.004, t = −2.095, p = 0.044). Table 2. Displays the full results of fixed effect models.

**Figure 5.**
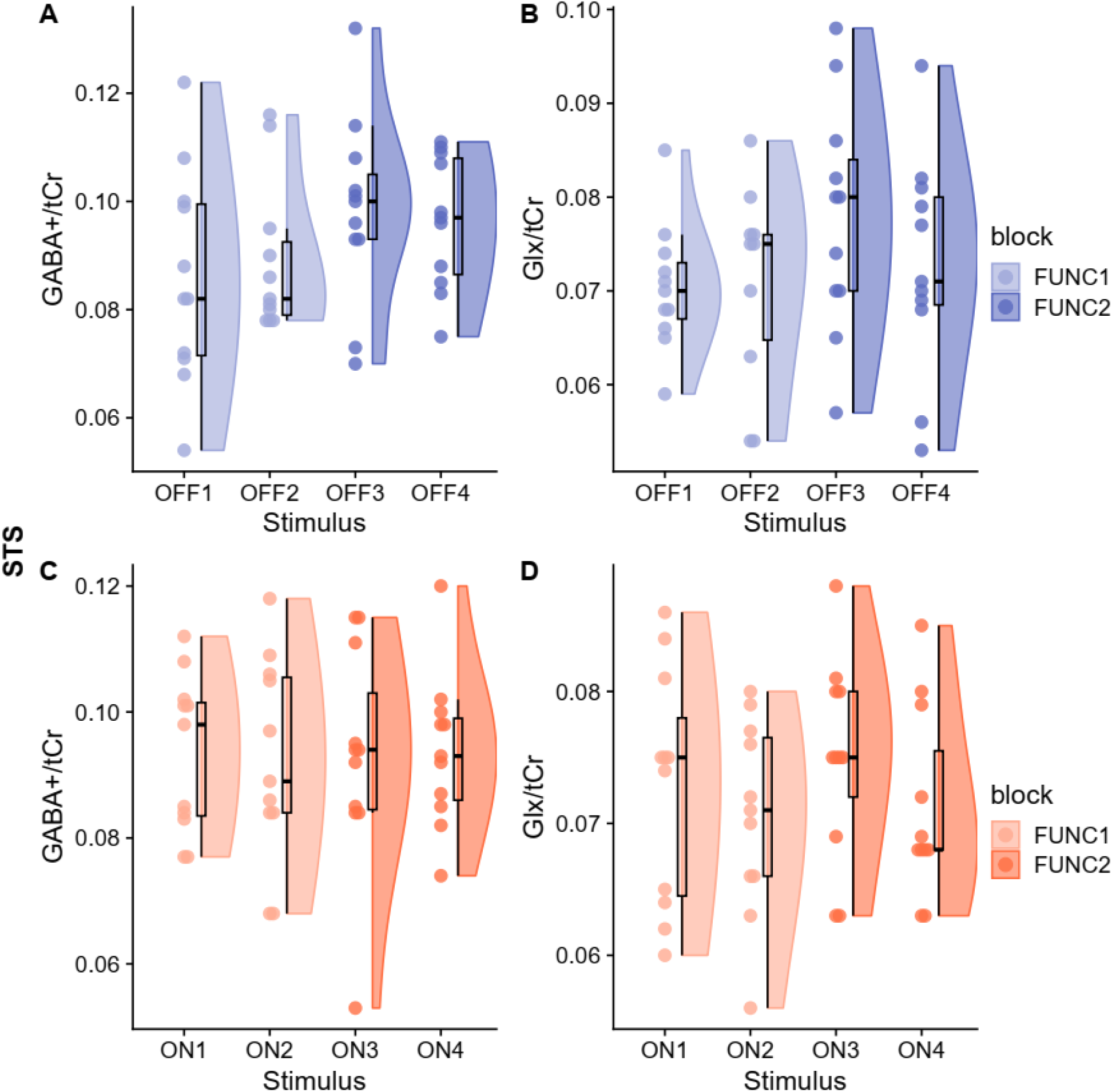
Raincloud plots of GABA+/tCr and Glx/tCr in STS during stimulus-ONs (24 sec stimulus-ONs of 3 sec movie clips of dynamic faces or dynamic objects) or stimulus-OFFs (16 sec rest [fixation cross]). ON_1-4_ are the stimulus-on TRs (regardless of stimulus type) within each functional block (ON1, ON2 for FUNC1 and ON3, ON4 for FUNC2). OFF_1-4_ are the stimulus-off TRs within each functional block (OFF1, OFF2 for FUNC1 and OFF3, OFF4 for FUNC2). P-values (<0.001***, <0.01**, and <0.05*) were obtained from a linear mixed model, and the full results are presented in Table 1.

**Figure 6.**
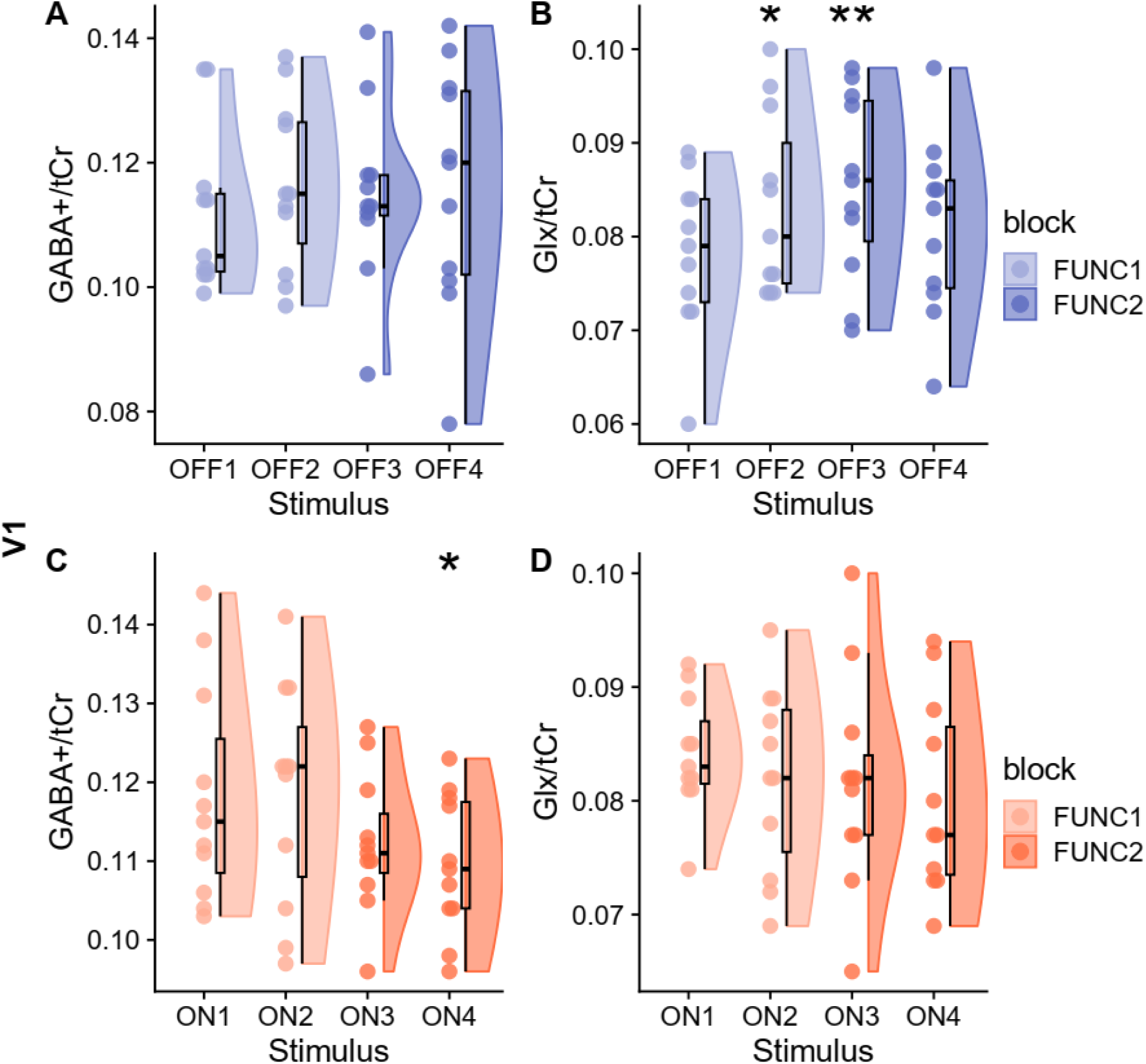
Raincloud plots of GABA+/tCr and Glx/tCr in V1 during stimulus-ONs (24 sec stimulus-ONs of 3 sec movie clips of dynamic faces or dynamic objects) or stimulus-OFFs (16 sec rest [fixation cross]). ON_1-4_ are the stimulus-on TRs (regardless of stimulus type) within each functional block (ON1, ON2 for FUNC1 and ON3, ON4 for FUNC2). OFF_1-4_ are the stimulus-off TRs within each functional block (OFF1, OFF2 for FUNC1 and OFF3, OFF4 for FUNC2). P-values (<0.001***, <0.01**, and <0.05*) were obtained from a linear mixed model, and the full results are presented in Table 1.

**Table 2.** Summary of linear mixed model fit for stimulus-ON and stimulus-OFF measurements of GABA+/tCR and Glx/tCr across functional stimulus types.

| Metabolite | Block | Predictors | Estimates($\beta$) | SE | df | t-value | P |
| --- | --- | --- | --- | --- | --- | --- | --- |
| V1 |  |  |  |  |  |  |  |
| GABA+/tCr | FUNC1 | OFF1 | Reference |  |  |  |  |
|  |  | OFF2 | 0.0046 | 0.0056 | 33 | 0.8230 | 0.417 |
|  | FUNC2 | OFF3 | 0.0032 | 0.0056 | 33 | 0.5650 | 0.576 |
|  |  | OFF4 | 0.0045 | 0.0056 | 33 | 0.8060 | 0.426 |
| GABA+/tCr | FUNC1 | ON1 | Reference |  |  |  |  |
|  |  | ON2 | 0.0003 | 0.0042 | 33 | 0.0650 | 0.948 |
|  | FUNC2 | ON3 | -0.0060 | 0.0042 | 33 | -1.4400 | 0.159 |
|  |  | ON4 | -0.0087 | 0.0042 | 33 | -2.0950 | 0.044* |
| Glx/tCr | FUNC1 | OFF1 | Reference |  |  |  |  |
|  |  | OFF2 | 0.0050 | 0.0021 | 33 | 2.3590 | 0.024* |
|  | FUNC2 | OFF3 | 0.0073 | 0.0021 | 33 | 3.4320 | 0.002** |
|  |  | OFF4 | 0.0028 | 0.0021 | 33 | 1.3300 | 0.193 |
| Glx/tCr | FUNC1 | ON1 | Reference |  |  |  |  |
|  |  | ON2 | -0.0022 | 0.0023 | 33 | -0.9610 | 0.343 |
|  | FUNC2 | ON3 | -0.0025 | 0.0023 | 33 | -1.0820 | 0.287 |
|  |  | ON4 | -0.0038 | 0.0023 | 33 | -1.6820 | 0.102 |
| STS |  |  |  |  |  |  |  |
| GABA+/tCr | FUNC1 | OFF1 | Reference |  |  |  |  |
|  |  | OFF2 | 0.0029 | 0.0065 | 33 | 0.4450 | 0.659 |
|  | FUNC2 | OFF3 | 0.0124 | 0.0065 | 33 | 1.8900 | 0.068 |
|  |  | OFF4 | 0.0103 | 0.0065 | 33 | 1.5710 | 0.126 |
| GABA+/tCr | FUNC1 | ON1 | Reference |  |  |  |  |
|  |  | ON2 | -0.0013 | 0.0044 | 33 | -0.2910 | 0.773 |
|  | FUNC2 | ON3 | -0.0005 | 0.0044 | 33 | -0.1250 | 0.902 |
|  |  | ON4 | 0.0003 | 0.0044 | 33 | 0.0620 | 0.951 |
| Glx/tCr | FUNC1 | OFF1 | Reference |  |  |  |  |
|  |  | OFF2 | 0.0008 | 0.0041 | 32.6 | 0.1850 | 0.854 |
|  | FUNC2 | OFF3 | 0.0075 | 0.0040 | 32.1 | 1.8900 | 0.068 |
|  |  | OFF4 | 0.0024 | 0.0040 | 32.1 | 0.5990 | 0.553 |
| Glx/tCr | FUNC1 | ON1 | Reference |  |  |  |  |
|  |  | ON2 | -0.0023 | 0.0026 | 33 | -0.8630 | 0.394 |
|  | FUNC2 | ON3 | 0.0021 | 0.0026 | 33 | 0.7940 | 0.433 |
|  |  | ON4 | -0.0016 | 0.0026 | 33 | -0.6210 | 0.539 |

### Sliding window analysis

The results obtained from the sliding window analysis indicate that there were fluctuating changes in Glx/tCr and GABA/tCr levels, with no obvious patterns observed irrespective of the type of stimulus presented in both V1 (Figure 7A, 7D) and STS (Figure 8A, 8D). However, it should be noted that the 95%CI for metabolite levels were relatively large when considering the metabolite levels based on the type of stimulus presented for both brain regions. This may be due to the low number of spectra available for certain stimulus types in each functional block.

**Figure 7.**
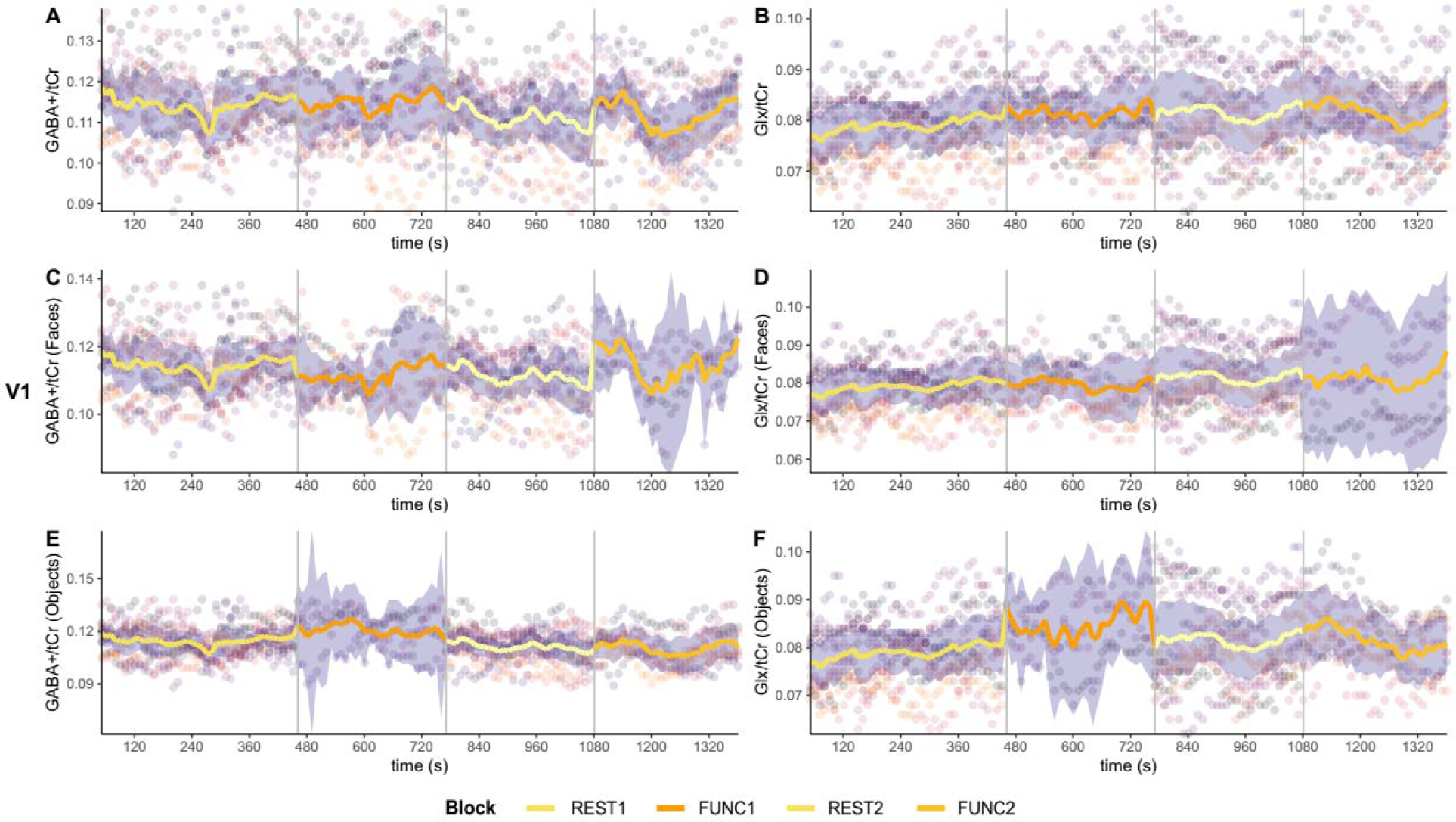
GABA+/tCr and Glx/tCr dynamics from sliding window analysis in V1 (window width = 50 transients). (A) GABA+/tCr and (B) Glx/tCr levels irrespective of stimulus type during FUNC1 and FUNC2. V1 GABA+/tCr levels in response to (C) faces or (E) objects stimuli during FUNC1 and FUNC2. V1 Glx/tCr levels in response to (D) faces or (F) objects stimuli during FUNC1 and FUNC2. The time in seconds represents the time at the centre of each window. The line fit represents the mean value of metabolite levels at each window, the purple ribbon indicates the 95% confidence interval (95% CI).

**Figure 8.**
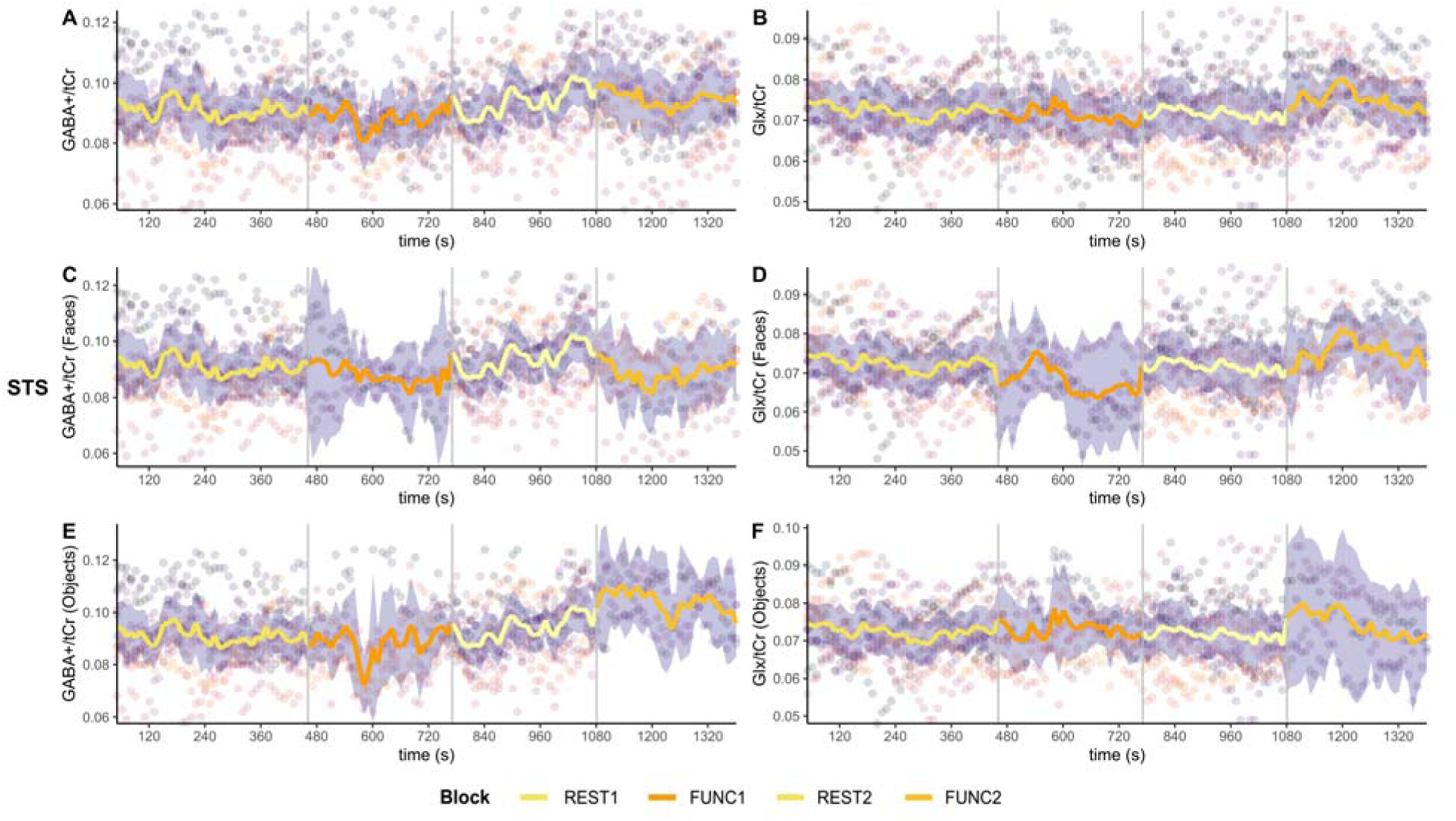
GABA+/tCr and Glx/tCr dynamics from sliding window analysis in STS (window width = 50 transients). (A) GABA+/tCr and (B) Glx/tCr levels irrespective of stimulus type during FUNC1 and FUNC2. STS GABA+/tCr levels in response to (C) faces or (E) objects stimuli during FUNC1 and FUNC2. STS Glx/tCr levels in response to (D) faces or (F) objects stimuli during FUNC1 and FUNC2. The time in seconds represents the time at the centre of each window. The line fit represents the mean value of metabolite levels at each window, the purple ribbon indicates the 95% confidence interval (95% CI).

The results of the linear mixed model analysis indicated that the stimulus type had a significant effect on V1 GABA+/tCr levels during FUNC1 (Table 3). Specifically, V1 GABA+/tCr increased for object stimuli (p < 0.001, β = 0.019, SE = 0.006) while decrease for faces stimuli (p < 0.001, β = −0.019, SE = 0.005) when compared to REST1.There was also a weak but significant interaction between time and faces stimuli presented in both FUNC1 (p = 0.001, β = 0.000026, SE = 0.000008) and FUNC2 (p = 0.03, β = −0.000022, SE = 0.03), while objects stimuli showed significant negative interaction with time during FUNC1 (p = 0.032, β = − 0.00002, SE = 0.00001). While both of the stimuli presented during FUNC1 significantly affect V1 Glx/tCr, the directionality was opposite with a positive relationship for objects (p = 0.029, β = 0.012, SE = 0.006) and a negative relationship for faces (p < 0.001, β = −0.015, SE = 0.004). Time also shown significant interaction with faces stimuli in FUNC1 (p = 0.028, β = −0.000014, SE = 0.000006) and REST2 (p = 0.028, β = −0.000014, SE = 0.000006).

**Table 3.**
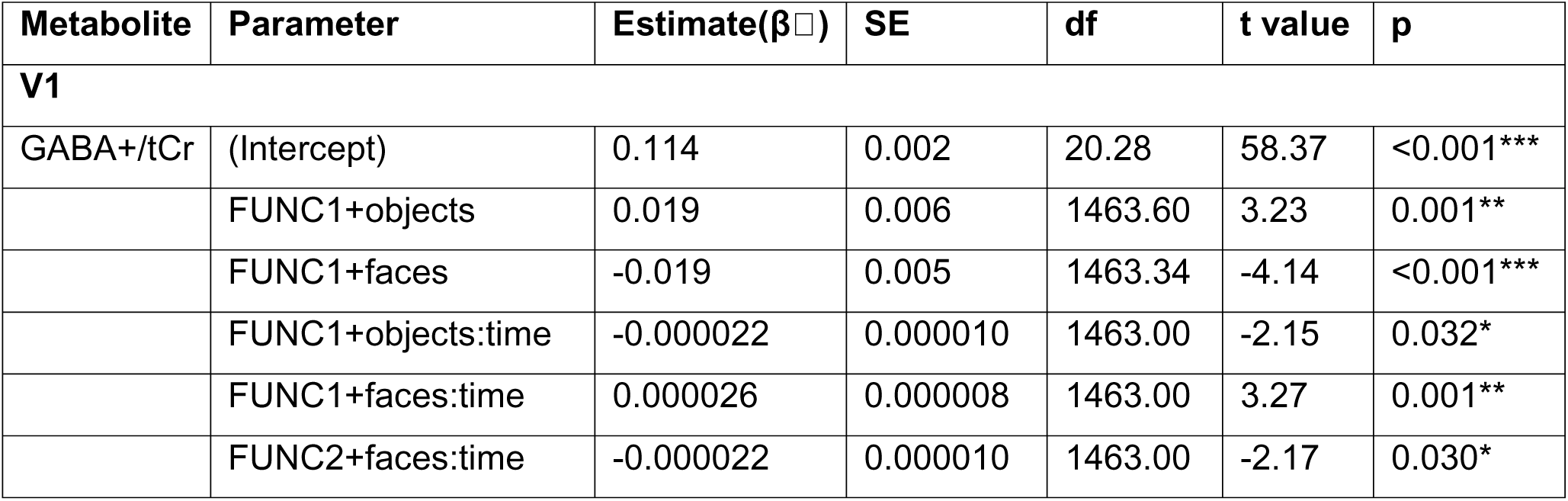
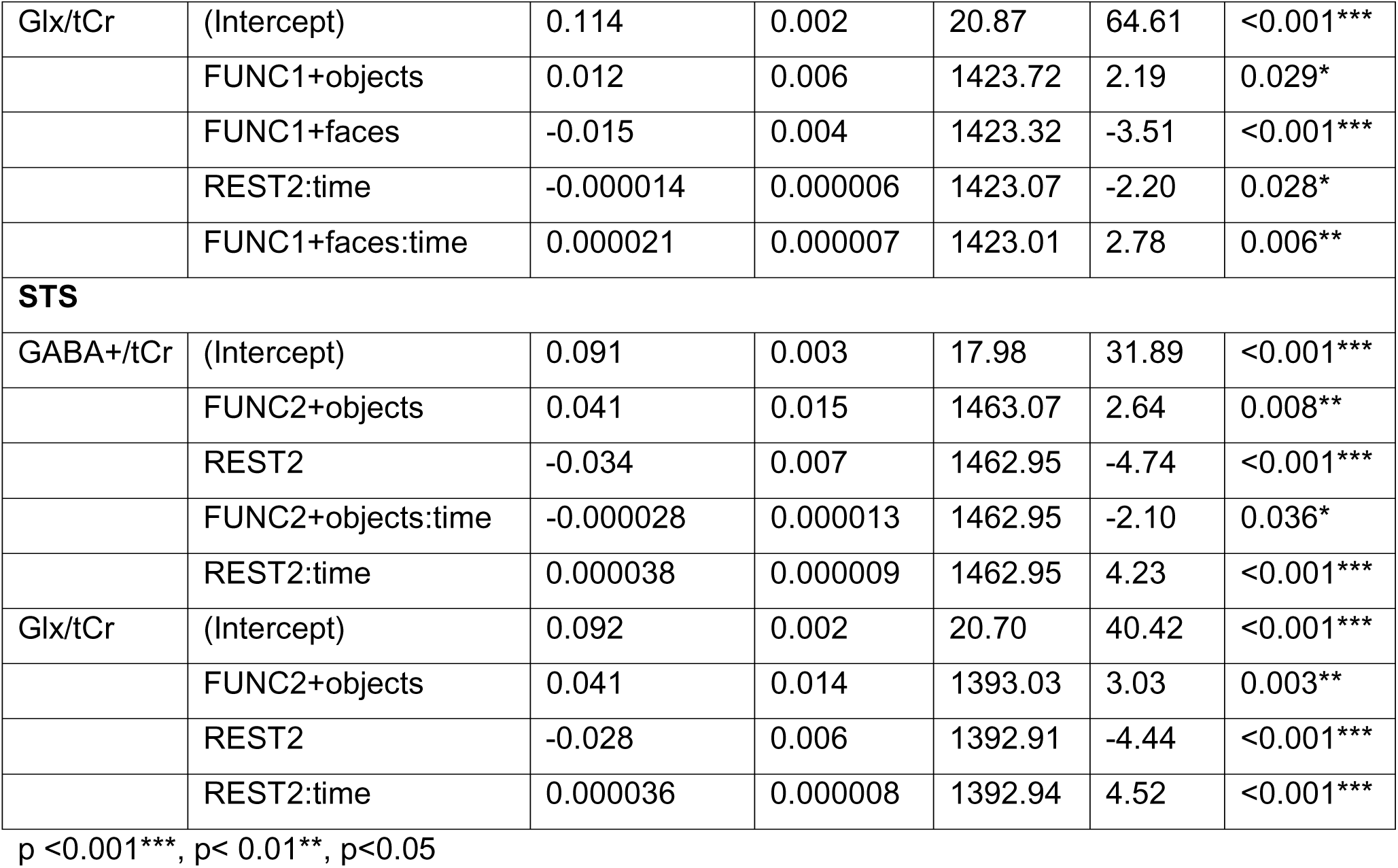
Summary of linear mixed model fit for sliding window analysis of GABA+/tCR and Glx/tCr in STS and V1. Only parameters with significant effect are shown here, for full parameters for each model please refer to Supplementary Material Table 1.

| Metabolite | Parameter | Estimate($\beta$) | SE | df | t value | p |
| --- | --- | --- | --- | --- | --- | --- |
| **V1** |  |  |  |  |  |  |
| GABA+/tCr | (Intercept) | 0.114 | 0.002 | 20.28 | 58.37 | <0.001*** |
|  | FUNC1+objects | 0.019 | 0.006 | 1463.60 | 3.23 | 0.001** |
|  | FUNC1+faces | -0.019 | 0.005 | 1463.34 | -4.14 | <0.001*** |
|  | FUNC1+objects:time | -0.000022 | 0.000010 | 1463.00 | -2.15 | 0.032* |
|  | FUNC1+faces:time | 0.000026 | 0.000008 | 1463.00 | 3.27 | 0.001** |
|  | FUNC2+faces:time | -0.000022 | 0.000010 | 1463.00 | -2.17 | 0.030* |
| Glx/tCr | (Intercept) | 0.114 | 0.002 | 20.87 | 64.61 | <0.001*** |
|  | FUNC1+objects | 0.012 | 0.006 | 1423.72 | 2.19 | 0.029* |
|  | FUNC1+faces | -0.015 | 0.004 | 1423.32 | -3.51 | <0.001*** |
|  | REST2:time | -0.000014 | 0.000006 | 1423.07 | -2.20 | 0.028* |
|  | FUNC1+faces:time | 0.000021 | 0.000007 | 1423.01 | 2.78 | 0.006** |
| **STS** |  |  |  |  |  |  |
| GABA+/tCr | (Intercept) | 0.091 | 0.003 | 17.98 | 31.89 | <0.001*** |
|  | FUNC2+objects | 0.041 | 0.015 | 1463.07 | 2.64 | 0.008** |
|  | REST2 | -0.034 | 0.007 | 1462.95 | -4.74 | <0.001*** |
|  | FUNC2+objects:time | -0.000028 | 0.000013 | 1462.95 | -2.10 | 0.036* |
|  | REST2:time | 0.000038 | 0.000009 | 1462.95 | 4.23 | <0.001*** |
| Glx/tCr | (Intercept) | 0.092 | 0.002 | 20.70 | 40.42 | <0.001*** |
|  | FUNC2+objects | 0.041 | 0.014 | 1393.03 | 3.03 | 0.003** |
|  | REST2 | -0.028 | 0.006 | 1392.91 | -4.44 | <0.001*** |
|  | REST2:time | 0.000036 | 0.000008 | 1392.94 | 4.52 | <0.001*** |

p <0.001\*\*\*, p< 0.01\*\*, p<0.05

On the other hand, linear mixed model showed no significant effect of social (faces) stimuli on STS GABA+/tCr and STS Glx/tCr. The results showed increase STS GABA+/tCr levels during objects stimulus presentation during FUNC2 compare to REST1 (p = 0.008, β = 0.041, SE = 0.015), while REST2 showed lower STS GABA+/tCr levels compared to REST1 (p < 0.036, β = −0.000028, SE = 0.000013). The objects stimuli during FUNC2 in STS also demonstrated a small negative interaction with time (p < 0.001, β = −0.015, SE = 0.004), whereas REST2 in STS showed a positive interaction with time (p < 0.001, β = 0.000038, SE = 0.000009). Similarly, the linear mixed model showed a significant positive relationship of objects stimulus presentation during FUNC2 on STS Glx/tCr (p = 0.003, β = −0.041, SE = 0.041), while REST2 demonstrated a significant lower STS Glx/tCr levels compared to REST1 (p < 0.001, β = − 0.028, SE = 0.006) with significant positive interaction with time (p < 0.001, β = 0.000036, SE = 0.000008).

### Changepoint analysis

Irrespective of stimulus types, the changepoint analysis demonstrated satisfactory fits with the provided models for both metabolites in V1 and STS (Rhat <1.1) (Figure 9). The exploratory changepoint analysis aimed to detect the initial shift in the trend of metabolites, demonstrated changes in both GABA+/tCr and Glx/tCr at the onset of the first rest block (REST1). The mean changepoint for V1 GABA+/tCr was determined to be 450.03 seconds, with a 95% CI (51.02, 1344.24), while for V1 Glx/tCr (Figure 9A), average changepoint was found to be 211 seconds, with a 95% CI (51.01, 769.05) (Figure 9B). The mean changepoint of 682.55 seconds, with a 95% CI [108.87, 1380.56], during the first functional block, was identified for STS GABA+/tCr (Figure 9A), and a changepoint of 967.23 seconds, with a 95% CI [182.95, 1380.91] during the REST2, was identified for STS Glx/tCr (Figure 9B). The relatively large 95% CI intervals indicate uncertainty in estimating the changepoint locations, as demonstrated by the posterior probability distributions presented in Figure 9.

**Figure 9.**
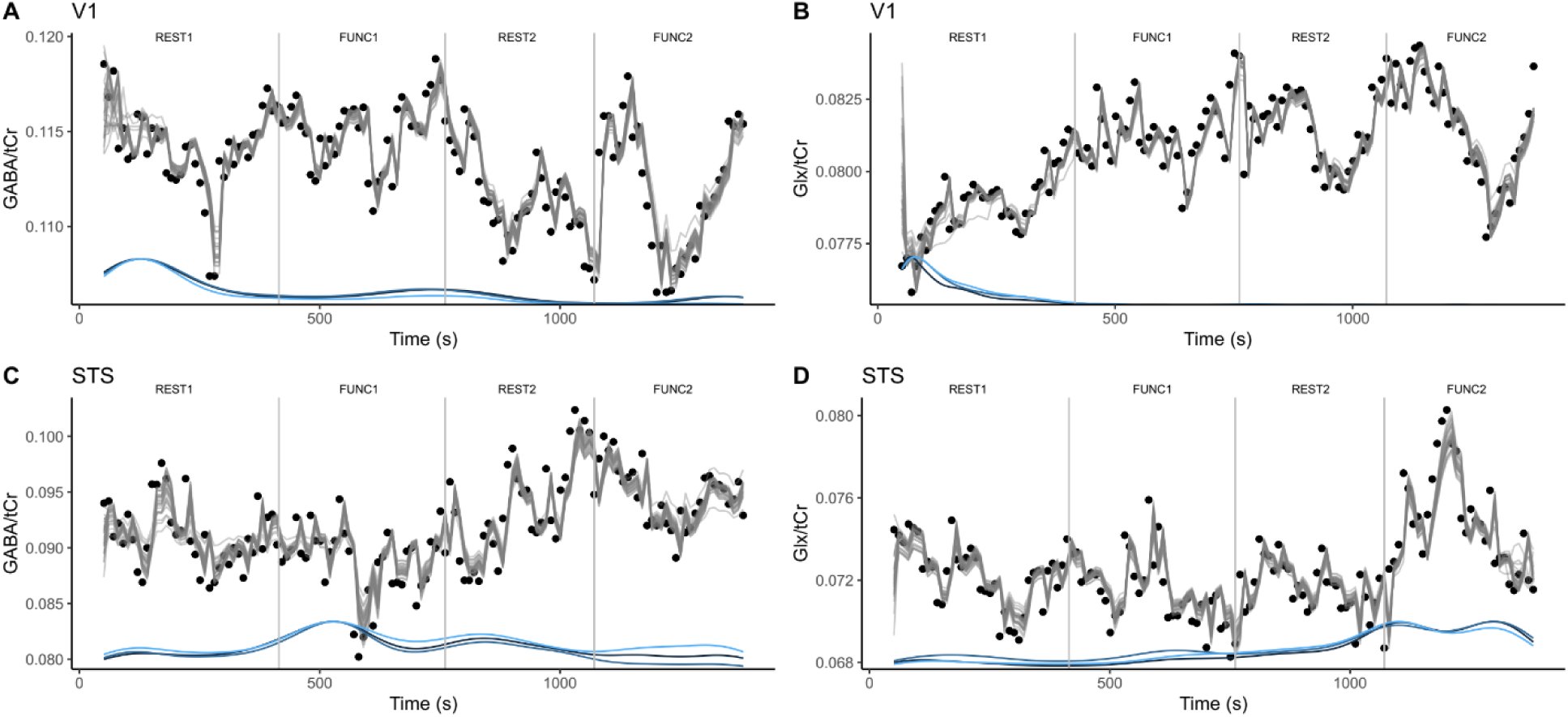
Results of changepoint analysis for (A, B) V1 and (C, D) STS regions. The gray/black line represents the posterior fit derived from the fitted models, the blue/black lines below each plot illustrates the posterior density of changepoints of each iteration chain at various time points. The height of the posterior density of changepoints signifies the potential values for changepoint locations, with the y-axis denoting the density (i.e., posterior probability) at each specific time point.

## Discussion

While studies suggest a connection between differences in social processes and alterations in glutamate and GABA function in neurodevelopmental disorders, the neurophysiology underlying this relationship remains largely unexplored. In this present study, we examined glutamate and GABA changes in response to social stimuli in non-clinical participants, using MEGA-PRESS functional MRS. Initial block-level analyses provided preliminary evidence that there was no association between the social processing task and changes in GABA+/tCr and Glx/tCr in both V1 and STS. Subsequent event-related analyses based on stimulus presentation events within each functional block (stimulus-ON or stimulus-OFF) indicated potential timing differences in how Glx/tCr and GABA+/tCr respond to stimulus presentation. Specifically, changes in V1 GABA+/tCr were significantly affected during the stimulus-ON in the last functional block, while changes in Glx/tCr were significantly impacted during stimulus-OFFs in both functional blocks (FUNC1 and FUNC2). Utilizing the sliding window analysis as an exploratory technique to examine metabolite changes at a higher temporal resolution, we demonstrated alterations in V1 Glx/tCr and Glx/tCr levels early in the first functional block while STS Glx/tCr and GABA+/tCr levels changes happened toward the last two functional blocks of REST2 and FUNC2.

When considering GABA and Glx levels using a block-type paradigm (RESTs vs FUNCs) we observed no significant changes between the rest and functional block for both V1 and STS. Subsequently, we also did not observe any significant differences between faces and objects stimuli. A possible explanation for this result is that a change in metabolite level is too marginal to be detected by fMRS with a typical number of transients (∼200 transients). While a block level analysis holds the advantage of the presumably summative effect of repeated stimuli within the same block, small metabolite responses might be averaged out with background changes (i.e., brain energetic consumption)^28^. Another possible explanation for the absence of significant changes for faces versus objects is that these stimuli are simply not salient enough to elicit a metabolite response with sufficient SNR, given the relatively short stimulation period given for each presentation of 24 seconds. A previous fMRI-fMRS study demonstrated a lack of significant metabolite level changes at low contrast but the response increased with increasing contrast levels^35^. Using sustained high-intensity visual stimulation, studies demonstrated small glutamate changes of only around 2%-4%^36–39^. These results suggest there is a possible sensitivity threshold for the functional stimulus to elicit a detectable response in fMRS. It is possible that this threshold will be largely dependent on the stimulus type and paradigm design as well ^19^.

We additionally investigated the metabolite change in an event-related design, separating transients into stimulus-ONs and stimulus-OFFs regardless of stimulus type to investigate potential lag of metabolite response with stimulus blocks; FUNC1 and FUNC2. For V1, we found a relative decrease in GABA+/tCr during the last stimulus-ON block, in which we hypothesised that might be due to accumulative change with continuous stimulus-ON TRs. This agrees with studies suggest that as stimulation continues, more GABA becomes bound to receptors, leading to linewidth broadening, and thus making it less detectable with MRS^40, 41^. The negative relationship of GABA with time might suggest that the evoked inhibitory tone does not completely return to baseline after stimulation, which has been previously suggested in recent animal optogenetic-fMRS studies^42^. Indeed, studies, including our recent meta-analysis^21^, suggest that GABA changes appear slowly.

With event-related analysis, we observed an increase in Glx/tCr relative to the start of stimulus-OFF (OFF1) in V1 toward the end of first functional block (OFF2) and during the first half of the second functional block (OFF3). This increase in Glx/tCr levels generally agrees with several studies showing an increase in glutamate in response to visual stimulation^19, 27, 38, 39, 43^. Interestingly, we found changes to be in stimulus-OFFs rather than stimulus-ON transients, suggesting a potential delay in the metabolic response of Glx/tCr in V1. This result is in agreement with previous studies that observed 3-6% glutamate increase after at least 16-20 seconds of stimulation as measured in a block paradigm^27^. Other studies which used event-related paradigms showed 9-12% glutamate changes after 300-1000 ms of stimulus onset^28, 44^. Here, we illustrated how an event-related paradigm might capture task-related transient metabolic changes that might be hidden by potential habituation and homeostatic regulation in a block paradigm.

In an attempt to establish the metabolite dynamics of both GABA+ and Glx further, the spectra were analysed using a sliding window method. This analysis method allows for metabolite quantification with higher temporal resolution, in our case with effective time course of 10 seconds (5 transients) with the metabolite levels averaged over 1 minute and 40 seconds (50 transients). Our sliding window analysis results demonstrated fluctuations of Glx/tCr and GABA+/tCr with no discernible pattern in both V1 and STS, irrespective of stimulus type.

In this study, both faces and objects stimuli demonstrated a significant effect on both V1 Glx/tCr and V1 GABA+/tCr during FUNC1, and the interaction terms suggest that changes are dependent on time. Despite the significant changes detected with event-related analysis during FUNC2, no significant changes in response to stimulus were observed during the FUNC2 with sliding window approach. This suggests that the changes during FUNC2 might be more abrupt or a quick change in trend, as the sliding window analysis approach could potentially smooth out such alterations in metabolite dynamics.

For STS, the linear mixed model did not indicate any significant effect of the social tasks (faces stimulus presentation on both STS Glx/tCr and STS GABA+/tCr). However, the results suggest a consistent effect of object stimulus presentation during FUNC2 on both Glx/tCr and GABA+/tCr, which is time-independent for GABA+/tCr. The results also showed a significant effect of the second rest block on both STS Glx/tCr and STS GABA+/tCr, with a significant interaction between REST2 and time of measurement suggesting time-dependence of the effect. It is possible that the significant effect of REST2, despite the absence of stimulus presentation, might be due to the nature of the sliding window analysis that includes the neighbour signal in other stimulus blocks (i.e., the previous FUNC1 block).

Changepoint analysis based on Bayesian inference approach reflected that GABA+/tCr and Glx/tCr levels acquired with sliding window analysis changes in a gradual fashion for stimuli in both V1 and STS, with a broad estimation of 95% CI of the change point. In comparison to STS, the change point in V1 is suggested to be appear early in the first rest block with changepoint of approximately 7 minutes for V1 GABA+/tCr and approximately 3 minutes V1 Glx/tCr. These results are in agreement with a recent study which investigated metabolite dynamics of GABA+ and Glx in visual cortex with high-resolution dynamic analysis and showed Glx and GABA drift despite the absence of stimulation^20^. Another explanation for the detected change point without the stimulus presentation might be due to the fixation cross used as the baseline subtly influencing the metabolite dynamics. It was previously demonstrated that different types of baseline tasks could affect glutamate levels^45^, or other brain processes captured by the Glx signal could be driving the Glx change and/or the detected Glx is influenced by Gln rather than glutamate ^20^.

On the other hand, the changepoint analysis identified the changepoint of STS GABA/tCtr at around 3 minutes after the start of FUNC1 (around 11 minutes from the start of the experiment), while the changepoint of ∼2 minutes after the start of REST was identified for Glx/tCr (around 17 minutes from the start of the experiment). These results agree with the linear mixed model analysis that found the significant effect of REST2 on STS Glx/tCr levels that depends on the time of measurement, suggesting glutamate might take roughly 2-3 minutes to reach its steady state through the synaptic reorganisation process after the stimulus onset^18, 36, 43^. The different changepoints identified between STS and V1 could be explained by timing differences between metabolite types and between brain regions as previous shown in several studies^19, 27^.

We did not observe significant changes in metabolites responses evoked by social task (i.e., faces stimuli) in the STS region with the sliding window analysis approach. While STS is a cortical region responsible for social perception from visual cues, in this current study only V1 showed significant changes to stimulation. These results are consistent regardless of analytical approach. Therefore, our findings likely reflect neurochemical responses to ‘basic’ visual processing rather than social processing *per se*. While previous studies have shown the sensitivity of fMRS in detecting glutamate changes that differ in the response to the presentation of either objects or abstract pictures^44^, this current study suggests that the Glx response is not specific to the type of visual stimulation. Another possible explanation for the changes observed in V1 but not STS is that V1 has higher proximity to the receiver coils, thus higher SNR and therefore increasing the statistical power to detect changes ^46^.

The current study does have some limitations. It should be noted that the sample size of this study is small thus the effects observed should be interpreted as a preliminary finding that requires future exploration with larger sample size. Another limitation of this study is that we cannot distinguish between intracellular and intercellular pools of metabolite due to the nature of fMRS. It is generally hypothesized that intracellular metabolite pools are involved in neurotransmission and extracellular pools are involved in maintaining excitation and inhibition balance^47^. Of course, it is well-known that GABA as measured through MEGA-PRESS J-difference editing contains macromolecules^48–50^, and we focused our approach on Glx (glutamate + glutamine) which likely reflects neurometabolism rather than neurotransmission. Furthermore, by using a J-difference editing technique, which relies on subtraction of edit-ON and edit-OFF transients, the temporal resolution is somewhat diminished^12, 49^. An additional limitation is that the social stimulus here was a passive viewing task, and as such was not inherently social. Perhaps future studies might benefit from presenting social stimulus that require some interaction with, e.g., identify smiling image as positive interaction, to better reflect social context.

Here, we used fMRS to investigate regional changes in glutamate and GABA levels during social cognition and functioning stimuli. The combined use of social stimulus and fMRS allowed for direct investigation of how regional metabolites, especially GABA and glutamate, change during social processing. The exact mechanism of how GABA and glutamate are involved in social functioning is, of course, complex and poorly understood. Further multimodal research with more participants is necessary in order to better understand the mechanisms involved in these social processing, which might be the key to understanding social impairments in clinical populations in the future.

## Author contributions

DP wrote the original draft of manuscript text and prepared figures. NP reviewed and editing. NP, DW and TF supervision the work and curate data. All authors reviewed the manuscript.

## Acknowledgements

The authors also acknowledge the facilities of Swinburne Neuroimaging (SNI) and its flagship funding from the National Imaging Facility (NIF) under the National Collaborative Researcher Infrastructure Strategy (NCRIS) implemented by the Australian Government

## Sponsors/ grant numbers

This study was funded by a Swinburne Neuroimaging Access and Support Scheme Enhanced Capability Grant (2002-TFDW). Dr Talitha Ford was funded by a Deakin University Dean’s Postdoctoral Research Fellowship.

## Abbreviations

AR: Autoregressive
E/I balance: Excitatory-inhibitory balance
fMRS: functional MRS
GABA: Gamma-aminobutyric acid
GABA+: Gamma-aminobutyric acid + macromolecules
Glx: Glutamate and glutamine
MM: Macromolecules
SNR: Signal to noise ratio
STS: Superior temporal sulcus
tCr: Creatine + phosphocreatine
V1: Visual cortex

## Notes

### Competing Interest Statement

The authors have declared no competing interest.

## References

1. Insel T, Cuthbert B, Garvey M, et al. Research Domain Criteria (RDoC): Toward a New Classification Framework for Research on Mental Disorders. Vol 167. Am Psychiatric Assoc; 2010:748–751.

2. Vrij A, Edward K, Roberts KP, Bull R. Detecting deceit via analysis of verbal and nonverbal behavior. Journal of Nonverbal behavior. 2000;24(4):239–263.

3. Puts NAJ, Wodka EL, Harris AD, et al. Reduced GABA and Altered Somatosensory Function in Children with Autism Spectrum Disorder. Autism Res. 2017;10(4):608–619. doi:10.1002/aur.1691

4. Sapey-Triomphe LA, Lamberton F, Sonié S, Mattout J, Schmitz C. Tactile hypersensitivity and GABA concentration in the sensorimotor cortex of adults with autism. Autism Research. 2019;12(4):562–575.

5. Hirata K, Egashira K, Harada K, et al. Differences in frontotemporal dysfunction during social and non-social cognition tasks between patients with autism spectrum disorder and schizophrenia. Sci Rep. 2018;8(1):3014. doi:10.1038/s41598-018-21379-w

6. Orhan F, Fatouros-Bergman H, Goiny M, et al. CSF GABA is reduced in first-episode psychosis and associates to symptom severity. Mol Psychiatry. 2018;23(5):1244–1250. doi:10.1038/mp.2017.25

7. Elsaid S, Rubin-Kahana DS, Kloiber S, Kennedy SH, Chavez S, Le Foll B. Neurochemical Alterations in Social Anxiety Disorder (SAD): A Systematic Review of Proton Magnetic Resonance Spectroscopic Studies. International Journal of Molecular Sciences. 2022;23(9):4754. doi:10.3390/ijms23094754

8. Horder J, Petrinovic MM, Mendez MA, et al. Glutamate and GABA in autism spectrum disorder—a translational magnetic resonance spectroscopy study in man and rodent models. Translational psychiatry. 2018;8(1):1–11.

9. Ford TC, Nibbs R, Crewther DP. Glutamate/GABA+ ratio is associated with the psychosocial domain of autistic and schizotypal traits. PloS one. 2017;12(7):e0181961.

10. Cochran DM, Sikoglu EM, Hodge SM, et al. Relationship among glutamine, γ-aminobutyric acid, and social cognition in autism spectrum disorders. Journal of child and adolescent psychopharmacology. 2015;25(4):314–322.

11. De Graaf RA. In vivo NMR spectroscopy: principles and techniques. Published online 2019.

12. Wilson M, Andronesi O, Barker PB, et al. Methodological consensus on clinical proton MRS of the brain: Review and recommendations. Magnetic Resonance in Medicine. 2019;82(2):527–550. doi:10.1002/mrm.27742

13. Öz G, Alger JR, Barker PB, et al. Clinical proton MR spectroscopy in central nervous system disorders. Radiology. 2014;270(3):658–679.

14. Rae CD. A Guide to the Metabolic Pathways and Function of Metabolites Observed in Human Brain 1H Magnetic Resonance Spectra. Neurochem Res. 2014;39(1):1–36. doi:10.1007/s11064-013-1199-5

15. Stagg CJ. Magnetic Resonance Spectroscopy as a tool to study the role of GABA in motor-cortical plasticity. NeuroImage. 2014;86:19–27. doi:10.1016/j.neuroimage.2013.01.009

16. Choi IY, Andronesi OC, Barker P, et al. Spectral Editing in 1H Magnetic Resonance Spectroscopy: Experts’ Consensus Recommendations. NMR Biomed. 2021;34(5):e4411. doi:10.1002/nbm.4411

17. Jelen LA, King S, Mullins PG, Stone JM. Beyond static measures: A review of functional magnetic resonance spectroscopy and its potential to investigate dynamic glutamatergic abnormalities in schizophrenia. J Psychopharmacol. 2018;32(5):497–508. doi:10.1177/0269881117747579

18. Stanley JA, Raz N. Functional Magnetic Resonance Spectroscopy: The “New” MRS for Cognitive Neuroscience and Psychiatry Research. Frontiers in Psychiatry. 2018;9:76. doi:10.3389/fpsyt.2018.00076

19. Mullins PG. Towards a theory of functional magnetic resonance spectroscopy (fMRS): A meta-analysis and discussion of using MRS to measure changes in neurotransmitters in real time. Scand J Psychol. 2018;59(1):91–103. doi:10.1111/sjop.12411

20. Rideaux R. Temporal Dynamics of GABA and Glx in the Visual Cortex. eNeuro. 2020;7(4). doi:10.1523/ENEURO.0082-20.2020

21. Pasanta D, He JL, Ford T, Oeltzschner G, Lythgoe DJ, Puts NA. Functional MRS studies of GABA and glutamate/Glx – A systematic review and meta-analysis. Neuroscience & Biobehavioral Reviews. 2023;144:104940. doi:10.1016/j.neubiorev.2022.104940

22. Pitcher D, Dilks DD, Saxe RR, Triantafyllou C, Kanwisher N. Differential selectivity for dynamic versus static information in face-selective cortical regions. Neuroimage. 2011;56(4):2356–2363. doi:10.1016/j.neuroimage.2011.03.067

23. Mescher M, Merkle H, Kirsch J, Garwood M, Gruetter R. Simultaneous in vivo spectral editing and water suppression. NMR Biomed. 1998;11(6):266–272. doi:10.1002/(sici)1099-1492(199810)11:6<266::aid-nbm530>3.0.co;2-j

24. Edden RAE, Puts NAJ, Harris AD, Barker PB, Evans CJ. Gannet: A batch-processing tool for the quantitative analysis of gamma-aminobutyric acid–edited MR spectroscopy spectra. Journal of Magnetic Resonance Imaging. 2014;40(6):1445–1452. doi:10.1002/jmri.24478

25. Mikkelsen M, Tapper S, Near J, Mostofsky SH, Puts NAJ, Edden RAE. Correcting frequency and phase offsets in MRS data using robust spectral registration. NMR in Biomedicine. 2020;33(10):e4368. doi:10.1002/nbm.4368

26. Near J, Edden R, Evans CJ, Paquin R, Harris A, Jezzard P. Frequency and phase drift correction of magnetic resonance spectroscopy data by spectral registration in the time domain. Magnetic Resonance in Medicine. 2015;73(1):44–50. doi:10.1002/mrm.25094

27. Betina Ip I, Berrington A, Hess AT, Parker AJ, Emir UE, Bridge H. Combined fMRI-MRS acquires simultaneous glutamate and BOLD-fMRI signals in the human brain. Neuroimage. 2017;155:113–119. doi:10.1016/j.neuroimage.2017.04.030

28. Apšvalka D, Gadie A, Clemence M, Mullins PG. Event-related dynamics of glutamate and BOLD effects measured using functional magnetic resonance spectroscopy (fMRS) at 3T in a repetition suppression paradigm. NeuroImage. 2015;118:292–300. doi:10.1016/j.neuroimage.2015.06.015

29. Leys C, Ley C, Klein O, Bernard P, Licata L. Detecting outliers: Do not use standard deviation around the mean, use absolute deviation around the median. Journal of Experimental Social Psychology. 2013;49(4):764–766. doi:10.1016/j.jesp.2013.03.013

30. Rousseeuw PJ, Croux C. Alternatives to the median absolute deviation. Journal of the American Statistical association. 1993;88(424):1273–1283.

31. Kuznetsova A, Brockhoff PB, Christensen RH. lmerTest package: tests in linear mixed effects models. Journal of statistical software. 2017;82:1–26.

32. Lindeløv JK. mcp: An R package for regression with multiple change points. Published online 2020.

33. van de Schoot R, Depaoli S, King R, et al. Bayesian statistics and modelling. Nat Rev Methods Primers. 2021;1(1):1–26. doi:10.1038/s43586-020-00001-2

34. Wagenmakers EJ, Marsman M, Jamil T, et al. Bayesian inference for psychology. Part I: Theoretical advantages and practical ramifications. Psychon Bull Rev. 2018;25(1):35–57. doi:10.3758/s13423-017-1343-3

35. Ip IB, Emir UE, Parker AJ, Campbell J, Bridge H. Comparison of Neurochemical and BOLD Signal Contrast Response Functions in the Human Visual Cortex. J Neurosci. 2019;39(40):7968–7975. doi:10.1523/JNEUROSCI.3021-18.2019

36. Mangia S, Tkác I, Gruetter R, Van de Moortele PF, Maraviglia B, Uğurbil K. Sustained neuronal activation raises oxidative metabolism to a new steady-state level: evidence from 1H NMR spectroscopy in the human visual cortex. Journal of cerebral blood flow and metabolism : official journal of the International Society of Cerebral Blood Flow and Metabolism. 2007;27(5):1055–1063. doi:10.1038/sj.jcbfm.9600401

37. Lin Y, Stephenson MC, Xin L, Napolitano A, Morris PG. Investigating the metabolic changes due to visual stimulation using functional proton magnetic resonance spectroscopy at 7LT. Journal of cerebral blood flow and metabolism : official journal of the International Society of Cerebral Blood Flow and Metabolism. 2012;32(8):1484–1495. doi:10.1038/jcbfm.2012.33

38. Bednařík P, Tkáč I, Giove F, et al. Neurochemical and BOLD responses during neuronal activation measured in the human visual cortex at 7 Tesla. Journal of cerebral blood flow and metabolism : official journal of the International Society of Cerebral Blood Flow and Metabolism. 2015;35(4):601–610. doi:10.1038/jcbfm.2014.233

39. Kurcyus K, Annac E, Hanning NM, et al. Opposite Dynamics of GABA and Glutamate Levels in the Occipital Cortex during Visual Processing. The Journal of neuroscience : the official journal of the Society for Neuroscience. 2018;38(46):9967–9976. doi:10.1523/JNEUROSCI.1214-18.2018

40. Jahnke W, Floersheim P, Ostermeier C, et al. NMR Reporter Screening for the Detection of High-Affinity Ligands. Angewandte Chemie International Edition. 2002;41(18):3420–3423. doi:10.1002/1521-3773(20020916)41:18<3420::AID-ANIE3420>3.0.CO;2-E

41. Rideaux R, Goncalves NR, Welchman AE. Mixed-polarity random-dot stereograms alter GABA and Glx concentration in the early visual cortex. Journal of neurophysiology. 2019;122(2):888–896. doi:10.1152/jn.00208.2019

42. Just N, Faber C. Probing activation-induced neurochemical changes using optogenetics combined with functional magnetic resonance spectroscopy: a feasibility study in the rat primary somatosensory cortex. Journal of Neurochemistry. 2019;150(4):402–419. doi:10.1111/jnc.14799

43. Schaller B, Mekle R, Xin L, Kunz N, Gruetter R. Net increase of lactate and glutamate concentration in activated human visual cortex detected with magnetic resonance spectroscopy at 7 tesla. Journal of neuroscience research. 2013;91(8):1076–1083. doi:10.1002/jnr.23194

44. Lally N, Mullins PG, Roberts MV, Price D, Gruber T, Haenschel C. Glutamatergic correlates of gamma-band oscillatory activity during cognition: a concurrent ER-MRS and EEG study. NeuroImage. 2014;85 Pt 2:823–833. doi:10.1016/j.neuroimage.2013.07.049

45. Lynn J, Woodcock EA, Anand C, Khatib D, Stanley JA. Differences in steady-state glutamate levels and variability between ‘non-task-active’ conditions: Evidence from 1H fMRS of the prefrontal cortex. Neuroimage. 2018;172:554–561. doi:10.1016/j.neuroimage.2018.01.069

46. Mikkelsen M, Loo RS, Puts NAJ, Edden RAE, Harris AD. Designing GABA-Edited Magnetic Resonance Spectroscopy Studies: Considerations of Scan Duration, Signal-To-Noise Ratio and Sample Size. J Neurosci Methods. 2018;303:86–94. doi:10.1016/j.jneumeth.2018.02.012

47. Belelli D, Harrison NL, Maguire J, Macdonald RL, Walker MC, Cope DW. Extrasynaptic GABAA receptors: form, pharmacology, and function. J Neurosci. 2009;29(41):12757–12763. doi:10.1523/JNEUROSCI.3340-09.2009

48. Mullins PG, McGonigle DJ, O’Gorman RL, et al. Current practice in the use of MEGA-PRESS spectroscopy for the detection of GABA. NeuroImage. 2014;86:43–52. doi:10.1016/j.neuroimage.2012.12.004

49. Peek AL, Rebbeck TJ, Leaver AM, et al. A comprehensive guide to MEGA-PRESS for GABA measurement. Analytical Biochemistry. 2023;669:115113. doi:10.1016/j.ab.2023.115113

50. Cudalbu C, Behar KL, Bhattacharyya PK, et al. Contribution of macromolecules to brain 1 H MR spectra: Experts’ consensus recommendations. NMR Biomed. 2021;34(5):e4393. doi:10.1002/nbm.4393

