## Supplementary material for "GABA and Glutamate response to social processing; a functional MRS study"

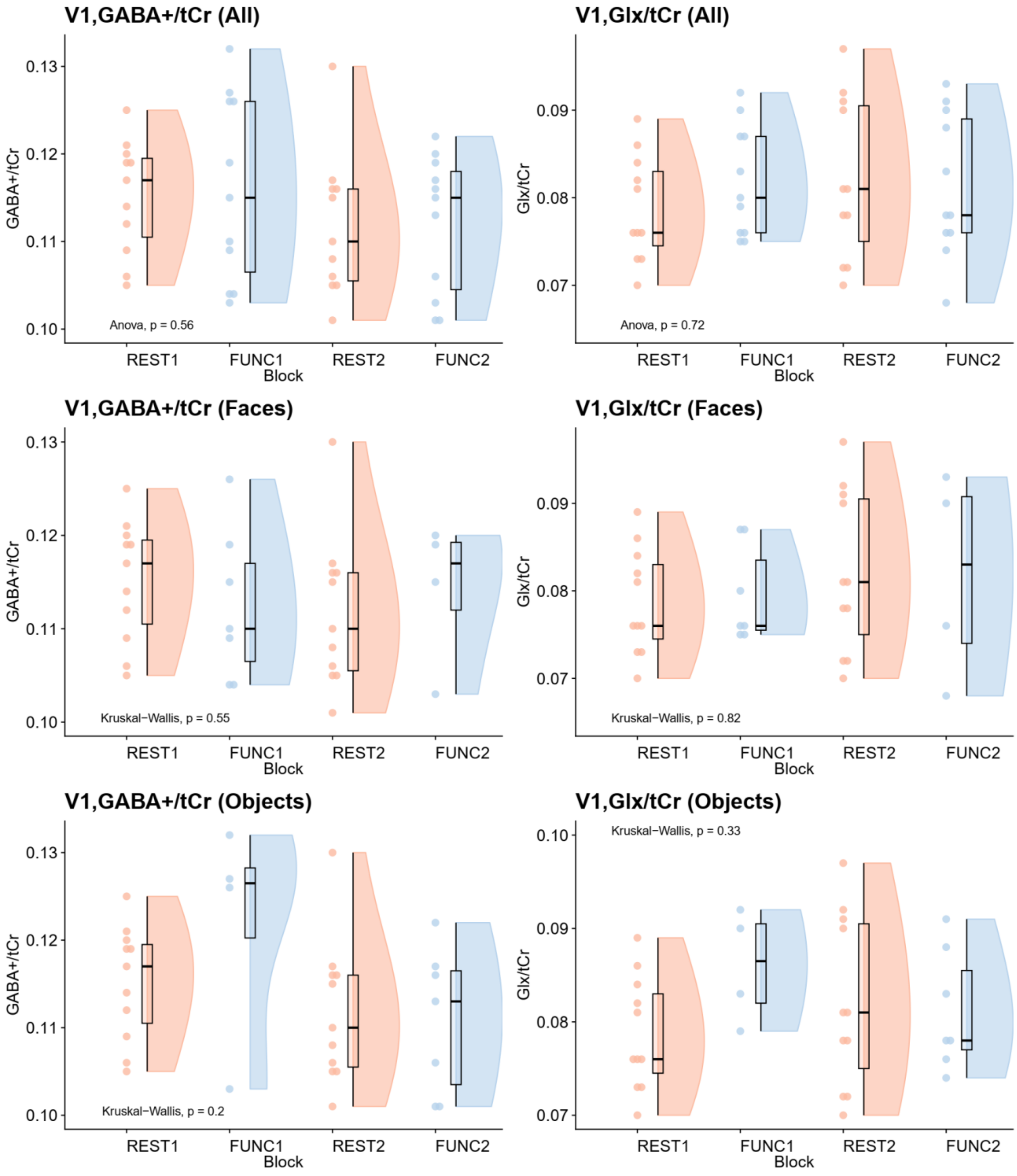


**Supplementary figure 1.** GABA+/tCr and Glx/tCr levels in V1 based on fMRS blocks for difference types of stimuli. All: both stimulus types (faces and objects) were included in the analysis, Faces: only metabolite levels during faces stimuli were included in the analysis; Objects: only metabolite levels during objects stimuli were included in the analysis. FUNC1, FUNC2: participants passively viewed social stimuli; REST1, REST2: participants passively viewed a fixation cross.

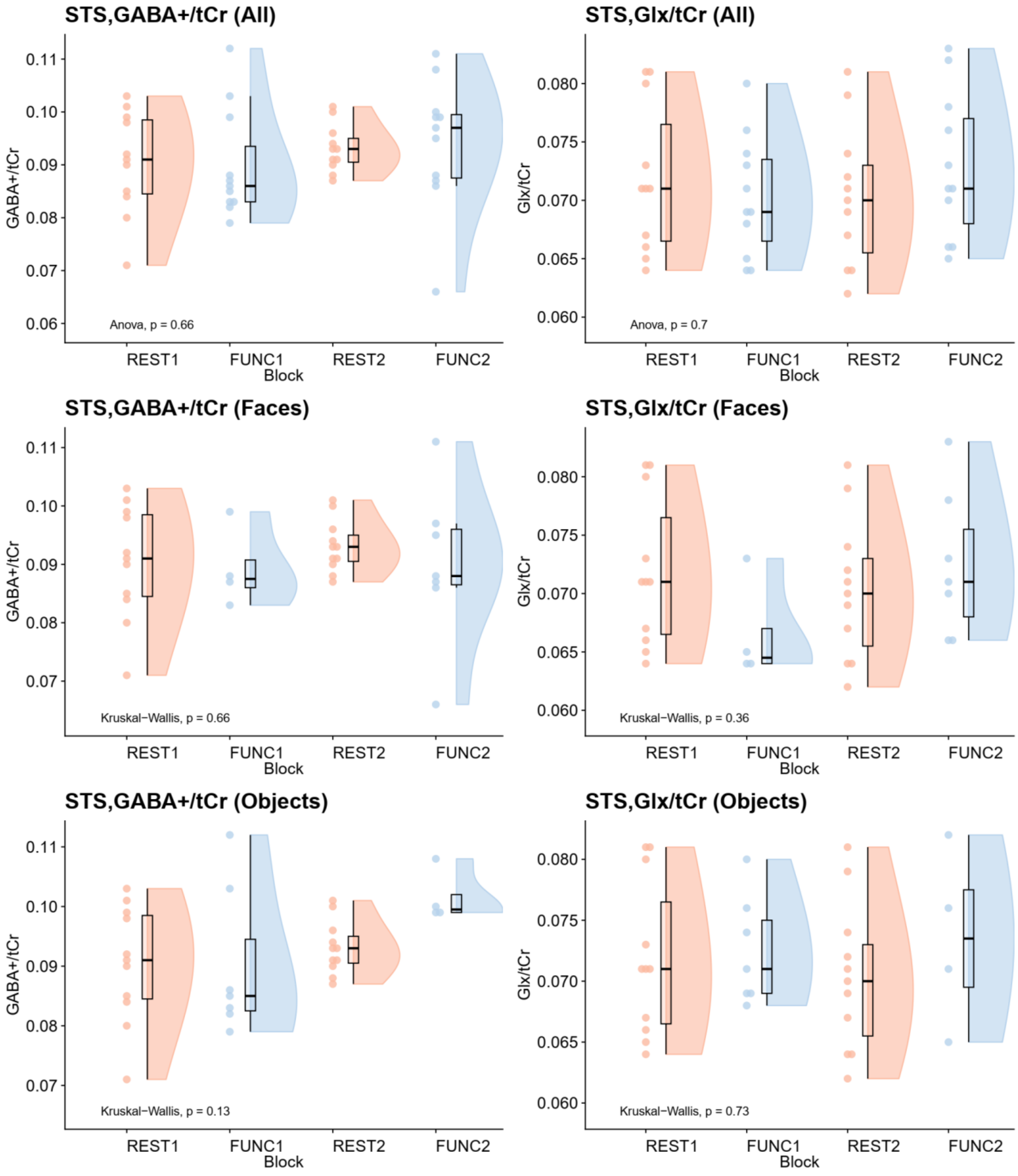


**Supplementary figure 2.** GABA+/tCr and Glx/tCr levels in STS and V1 based on fMRS blocks across both stimulus types. FUNC1, FUNC2: participants passively viewed social stimuli; REST1, REST2: participants passively viewed fixation cross.

Supplementary Material Table 1. Summary of linear mixed model fit for sliding window analysis of GABA+/tCR and Glx/tCr in STS and V1. Only parameters with significant effect are shown here, for full parameters for each model please refer to Table 1 in Supplementary Material.

| Metabolite | Parameter | Estimate | SE | df | t value | p |
|---|---|---|---|---|---|---|
| V1 | | | | | | |
| GABA | (Intercept) | 0.114 | 0.002 | 20.28 | 58.37 | <0.001*** |
| | FUNC1+objects | 0.019 | 0.006 | 1463.60 | 3.23 | 0.001** |
| | FUNC2+objects | -0.002 | 0.009 | 1463.09 | -0.19 | 0.846 |
| | REST2 | 0.004 | 0.005 | 1463.00 | 0.81 | 0.416 |
| | FUNC1+faces | -0.019 | 0.005 | 1463.34 | -4.14 | <0.001*** |
| | FUNC2+faces | 0.022 | 0.012 | 1463.16 | 1.93 | 0.054 |
| | time | 0.000003 | 0.000004 | 1463.00 | 0.72 | 0.473 |
| | FUNC1+objects:time | -0.000022 | 0.000010 | 1463.00 | -2.15 | 0.032* |
| | FUNC2+objects:time | -0.000003 | 0.000008 | 1463.00 | -0.42 | 0.672 |
| | REST2:time | -0.000010 | 0.000007 | 1463.00 | -1.41 | 0.160 |
| | FUNC1+faces:time | 0.000026 | 0.000008 | 1463.00 | 3.27 | 0.001** |
| | FUNC2+faces:time | -0.000022 | 0.000010 | 1463.00 | -2.17 | 0.030* |
| Glx | (Intercept) | 0.114 | 0.002 | 20.87 | 64.61 | <0.001*** |
| | FUNC1+objects | 0.012 | 0.006 | 1423.72 | 2.19 | 0.029* |
| | FUNC2+objects | 0.000 | 0.008 | 1423.08 | 0.04 | 0.968 |
| | REST2 | 0.010 | 0.005 | 1423.09 | 1.93 | 0.054 |
| | FUNC1+faces | -0.015 | 0.004 | 1423.32 | -3.51 | <0.001*** |
| | FUNC2+faces | 0.013 | 0.011 | 1423.19 | 1.22 | 0.223 |
| | time | 0.000000 | 0.000003 | 1423.03 | 0.11 | 0.912 |
| | FUNC1+objects:time | -0.000012 | 0.000010 | 1423.07 | -1.23 | 0.219 |
| | FUNC2+objects:time | -0.000003 | 0.000007 | 1423.00 | -0.42 | 0.672 |
| | REST2:time | -0.000014 | 0.000006 | 1423.07 | -2.20 | 0.028* |
| | FUNC1+faces:time | 0.000021 | 0.000007 | 1423.01 | 2.78 | 0.006** |
| | FUNC2+faces:time | -0.000012 | 0.000009 | 1423.02 | -1.23 | 0.217 |
| STS | | | | | | |
| GABA | (Intercept) | 0.091 | 0.003 | 17.98 | 31.89 | <0.001*** |
| | FUNC1+objects | -0.008 | 0.006 | 1463.22 | -1.40 | 0.161 |
| | FUNC2+objects | 0.041 | 0.015 | 1463.07 | 2.64 | 0.008** |
| | REST2 | -0.034 | 0.007 | 1462.95 | -4.74 | <0.001*** |
| | FUNC1+faces | 0.006 | 0.008 | 1463.43 | 0.70 | 0.483 |
| | FUNC2+faces | 0.003 | 0.012 | 1463.02 | 0.28 | 0.778 |
| | time | 0.000003 | 0.000005 | 1462.95 | 0.54 | 0.590 |
| | FUNC1+objects:time | 0.000018 | 0.000011 | 1462.95 | 1.69 | 0.092 |
| | FUNC2+objects:time | -0.000028 | 0.000013 | 1462.95 | -2.10 | 0.036* |
| | REST2:time | 0.000038 | 0.000009 | 1462.95 | 4.23 | <0.001*** |

| | | | | | | |
|---|---|---|---|---|---|---|
| | FUNC1+faces:time | -0.000023 | 0.000013 | 1462.95 | -1.71 | 0.088 |
| | FUNC2+faces:time | -0.000004 | 0.000011 | 1462.95 | -0.39 | 0.696 |
| Glx | (Intercept) | 0.092 | 0.002 | 20.70 | 40.42 | <0.001*** |
| | FUNC1+objects | -0.008 | 0.005 | 1393.31 | -1.47 | 0.142 |
| | FUNC2+objects | 0.041 | 0.014 | 1393.03 | 3.03 | 0.003** |
| | REST2 | -0.028 | 0.006 | 1392.91 | -4.44 | <0.001*** |
| | FUNC1+faces | 0.008 | 0.007 | 1393.47 | 1.15 | 0.252 |
| | FUNC2+faces | -0.004 | 0.011 | 1393.21 | -0.35 | 0.724 |
| | time | -0.000004 | 0.000004 | 1393.02 | -0.81 | 0.419 |
| | FUNC1+objects:time | 0.000016 | 0.000010 | 1392.94 | 1.68 | 0.093 |
| | FUNC2+objects:time | -0.000023 | 0.000012 | 1392.90 | -1.92 | 0.055 |
| | REST2:time | 0.000036 | 0.000008 | 1392.94 | 4.52 | <0.001*** |
| | FUNC1+faces:time | -0.000022 | 0.000012 | 1392.90 | -1.84 | 0.067 |
| | FUNC2+faces:time | 0.000006 | 0.000010 | 1393.02 | 0.62 | 0.535 |

p <0.001***, p< 0.01**, p<0.05
